# PhagePickr: A bacteria-centric computational tool for designing resistance-aware phage cocktails

**DOI:** 10.64898/2026.03.23.713575

**Authors:** Alessandro Oneto, Kenichi W. Okamoto

## Abstract

As antibiotic resistance poses a major threat to global health, phage therapy offers an alternative to antibiotic treatments in the face of multidrug-resistant bacteria. However, host resistance to phages is also well-documented. Current computational tools for phage cocktail design do not explicitly address the evolution of phage resistance, let alone through the profiling of bacterial receptors whose variability drives much of phage resistance.

We introduce PhagePickr, a computational pipeline for the automated design of phage cocktails intended to reduce the risk of host resistance. Unlike other tools, PhagePickr selects phages based on bacterial surface receptor similarity and prioritizes genomic diversity with the aim of reducing cross-resistance. The tool uses receptor annotations to identify phenotypically similar hosts via a nearest-neighbor algorithm and uses multiple sequence alignments to ensure genomic diversity in the final cocktail.

We evaluated the utility of PhagePickr on ESKAPE pathogens and two understudied bacterial species. The cocktails included candidate phages selected using bacterial receptor similarity and phage genomic divergence, comprising both lytic phages with confirmed therapeutic potential and novel candidates from similar species. We demonstrate the tool’s utility in generating cocktails and its capacity to scale as current databases are updated. PhagePickr provides a novel bacteria-centric framework for resistance-aware cocktail design by exploring shared phenotypes.

**Author Summary:** We present PhagePickr, a novel computational tool to design bacteriophage cocktails against pathogenic bacteria. Antibiotic resistance poses a major threat to global health, and phage therapy, the use of viruses that kill bacteria, is a promising alternative treatment. However, bacteria are also under immense selective pressure to develop resistance to phages, and existing tools for automated cocktail design have yet to address this challenge. Our tool aims to reduce the risk of resistance through two steps: it uses bacterial receptor configurations as predictors of phage infection and constructs a cocktail that maximizes phage genomic diversity, under the hypothesis that selected phages may target different receptors and make simultaneous resistance harder to evolve. We demonstrated the utility of PhagePickr on two understudied species and the ESKAPE pathogens, a group of multidrug-resistant bacteria responsible for the majority of deaths associated with antibiotic resistance worldwide. The tool identified both well-characterized therapeutic phages and novel candidates, and is designed to scale as databases expand. Our approach represents a key step toward the rational design of resistance-informed phage cocktails.

## Introduction

Since their discovery, antibiotics have been widely used to treat bacterial infections. The emergence of antibiotic-resistant bacteria has initiated an arms race which antibiotics are currently losing. The discovery rate of new antibiotics has stagnated, possibly rendering antibiotic therapy ineffective against future infections (1). Further, the impacts of antibiotic-resistant bacteria on mortality are significant and pose a substantial threat to public health, causing an estimated 1.27 million deaths worldwide in 2019 alone (2,3).

To alleviate this public health burden, alternative treatments to antibiotics are needed. Bacteriophages (phages), viruses that infect bacteria, represent a potential option. These were used as therapeutic agents until they were eclipsed by the discovery and subsequent mass production of antibiotics (4). Nonetheless, the study of phages and phage-host interactions remains relevant due to their many possible applications in human health, agriculture, and food safety (5).

Phage therapy entails the use of phages to treat bacterial infections and has been proposed as a promising alternative or supplement to antibiotics against multidrug-resistant bacteria, such as *Clostridioides difficile* (6). Current evidence indicates efficient phage therapies in mice against *Klebsiella pneumoniae* (7) and *Shigella sonnei* (8). This therapeutic potential also extends to human trials for infections caused by important pathogens such as *Pseudomonas aeruginosa* (9) and *Staphylococcus aureus* (10–12) often without serious adverse effects on patients. However, the clinical application of phages is constrained by their intricate interactions with bacteria.

Unlike antibiotics, phages exhibit remarkable selectivity and specificity towards their bacterial hosts (4). A key driver of host-specificity is their binding to surface receptors in bacteria (13). For instance, phages can bind to virulence factors such as the Vi capsular antigen in *Salmonella enterica* (14) and the capsular polysaccharide in *Escherichia coli* (15). Yet even though they often have a confined host range, several phages including multiple bacterial genera in their host range have also been reported (16,17).

These broad-host-range phages offer many advantages as therapeutic agents, since they can target co-infections involving multiple bacterial species, infect diverse strains within a species, and most importantly, minimize phage-host mismatch to maximize binding efficacy (18). Although phages with narrow host ranges constitute the majority (19), the recent isolation of broad-host-range phages from different host receptors suggests they may be more prevalent than expected (18).

Host range is governed in part by phage morphology, and in particular by the compatibility between phage receptor-binding proteins (RBPs) and bacterial surface receptors (20,21). Many efforts have been directed at the study of RBPs, including their identification, functionality, and their potential to predict hosts (16,22). While most research is focused on RBPs, the role of surface receptors in host recognition has been only recently emphasized (23). Hence, the interplay between RBPs and bacterial receptors not only determines host range and successful phage binding (16) but also dictates the efficiency of phage therapy.

The receptor binding mechanism presents two main obstacles for phage therapy. First, it complicates phage selection by causing strict host specificity. Second, it provides a target for bacterial resistance to evolve. Like antibiotic resistance, bacteria can modify surface receptors or hide these under barriers, thereby blocking phage infection (13,17). This emergence of phage resistance in bacteria represents an important challenge to phage therapy (24,25).

To counter phage resistance, previous studies have extensively discussed phage cocktails, combinations of therapeutic phages with superior efficacy against pathogens like *Pseudomonas aeruginosa* (26) and *Salmonella* Typhimurium (27) over individual phages. Several examples of combination therapy exist in medicine, such as Highly Active Antiretroviral Therapy (HAART) for HIV or antibiotic cocktails for tuberculosis, which are designed to overcome resistance. Increasing phage diversity, as in cocktails, can prevent the development of phage resistance as bacteria may struggle to become resistant to multiple phages simultaneously (28).

A key advantage of phage therapy is its potential synergy with antibiotics, offering increased selective pressure to circumvent resistance in the host (29). The evolution of phage resistance in bacteria can even re-sensitize bacteria to antibiotics (9,25) by downregulating surface proteins involved in drug resistance (30,31). This highlights the value of cocktails designed to target multiple bacterial surface receptors.

Several bioinformatics tools predict phage-host interactions (21,32,33) or identify bacterial hosts from phage genomes (34,35). For automated cocktail design, the R package PhageCocktail is a key resource that designs effective cocktails via phage infection networks (36). However, none of the current approaches explicitly address the crucial challenge of phage resistance.

Here, we present PhagePickr, a computational tool for the systematic design of resistance-aware phage cocktails. To address resistance, PhagePickr employs a bacteria-centric receptor profiling approach. The tool selects phages based on bacterial surface receptor similarity, with the aim of identifying phages that target phenotypically similar hosts through different receptors and reduce the risk of bacterial resistance. Our objective is to facilitate the in silico cocktail design process, thereby contributing to the advancement of targeted phage therapy designed to minimize resistance evolution.

Unlike traditional phage cocktail design, which requires receptor characterization to target multiple pathways (25), PhagePickr computationally infers receptor-targeting potential from genomic data to select phages from bacteria with similar receptor profiles while maximizing genomic divergence among candidates (Fig 1). This approach is designed to reduce the risk of cross-resistance under the hypothesis that divergent phages use different infection strategies and could require bacteria to acquire multiple independent resistance mutations (37). PhagePickr thus relies on two key assumptions: similar receptor profiles predict phage cross-infectivity (or at least identify targets for genetic engineering or directed evolution) (38–40); and phage genomic divergence implies functional diversity (41). This functional diversity could increase the likelihood that phages employ different surface receptors to infect their host, thereby reducing the risk of phage resistance in bacteria.

**Fig 1.**
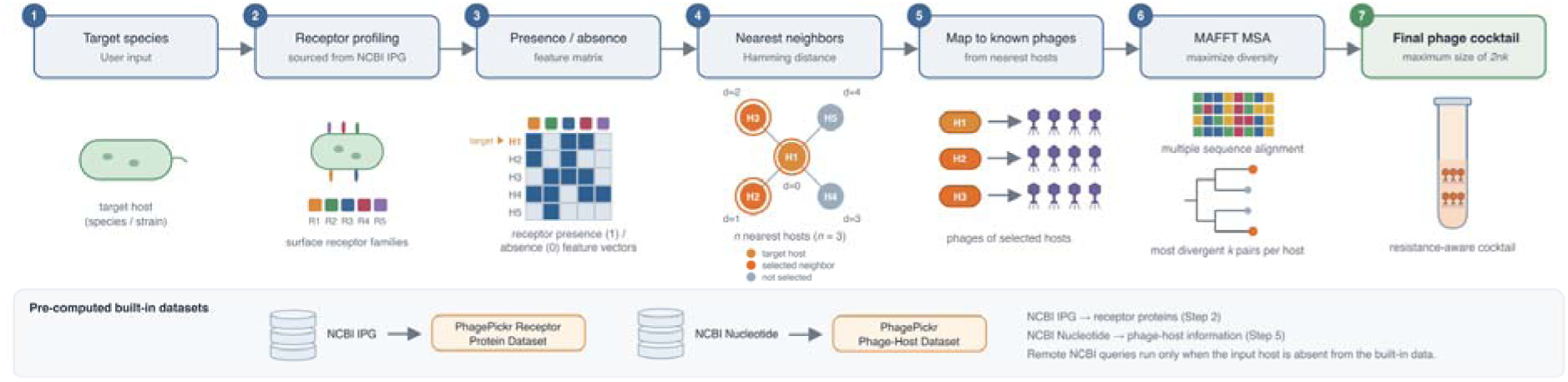
Overview of the PhagePickr workflow for resistance-aware cocktail design. The pipeline integrates bacterial receptor data (NCBI IPG) and phage-host data (NCBI Nucleotide) as built-in datasets, with remote NCBI queries only when the target is absent from them. Receptor proteins of the target species (Step 2) are encoded as a presence/absence vector over receptor features (Step 3). The target is then compared with all hosts in the built-in dataset, and the *n* closest are retained (Step 4), with the target included by default, contributing its known phages. Phages of the *n* selected hosts (Step 5) are aligned per host with MAFFT, and the *k* most divergent phage pairs from each alignment are selected (Step 6), generating a cocktail of at most 2*nk* phages (Step 7). The number of receptor features that differ from the target is represented by *d*. By default, PhagePickr computes the normalized Hamming distance, which gives the same neighbor ordering. The schematic shows an example with default parameters *n* = 3 and *k* = 1.

### Design and Implementation

PhagePickr identifies species with similar receptor configurations to select candidate phages that may infect the target. The aim of reducing resistance is motivated by evidence that bacteria under strong phage selection can alter the expression of targeted receptors, exposing vulnerabilities that could be exploited by antibiotics (9,42) or increasing susceptibility to other phage phenotypes (43,44). Although RBP stringency and specificity limit cross-infection potential (16), shared receptor configurations may increase the likelihood that some phages are effective against the target.

### Receptor Feature Matrix

Bacterial receptor and phage-host data of 200 pathogenic species were retrieved from the National Center for Biotechnology Information (NCBI)’s Identical Protein Groups (IPG) and Nucleotide databases (45) to construct two internal datasets and minimize queries at runtime: a record of receptor proteins per species, and a file recording hosts and their phage infectors. Both built-in datasets are provided as standalone resources on GitHub: the phage-host records (phagedicts.json) and the per-species receptor protein titles (receptor_data.json) from which the feature matrix is constructed. This enables use cases beyond phage cocktail design, such as feature extraction for machine learning models or the assembly of phage-bacteria interaction networks. While these datasets reflect the NCBI databases at the time of commit, the provided scripts allow users to generate the latest version locally.

The built-in receptor dataset is used to construct a binary presence/absence matrix 0,1 over *S*=200 pathogenic species and *P*=2,951 receptor-associated protein features. The species were obtained from a curated list of 201 pathogens (uniquepat.txt); one listed species returned no records and was excluded. For each species, candidate proteins were retrieved from NCBI’s IPG database using the Entrez query “ <species>[ORGN] AND receptor[All fields]”, run once per species (using both “esearch” and “esummary”, in batches of 8,000). The [ORGN] field fetches the organism following NCBI taxonomy, and the IPG database merges redundant protein records into one. Each feature is the record title. The matrix column space is composed of the aggregated titles across the 200 species, from which 139 empty or uninformative titles (e.g., “unnamed protein”) were dropped, leaving *P*=2,951 features. *A_s,p_* = 1 if species *s* contains protein *p*, and 0 otherwise. Of the 200 2,951 590,200 cells, 15,145 are populated (density 2.57%, mean of 76 features per species). Encoding the presence/absence of features rather than the amino acid or nucleotide sequence is robust to protein sequence variation and enables the efficient comparison of the receptor-protein phenotype across multiple species.

The feature space was obtained via the term “receptor[All fields]”, instead of a combination of specific receptor-related keywords. Because PhagePickr aims to generalize across diverse pathogens, we believe any additional term would further bias the feature space towards a particular clade and would impede adequate inter-species comparison. “receptor[All fields]” is clade-neutral, which minimizes bias at the cost of some annotation noise. It is also broader than a title match, given that only 69.0% (2,035/2,951) of feature columns contain the word “receptor”, while the remaining records reference the term in other fields. This naturally includes surface receptor proteins (e.g. OmpA, Tsx), but also some off-target proteins (e.g. kinases and chemotaxis proteins). The choice of term might exclude some well-characterized phage receptors, such as OmpC and OmpF in *E. coli*, which are annotated as porins instead of receptors and are thus absent from the matrix. Both excluding actual receptors and including off-target proteins introduce noise into the Hamming distance between pathogenic species rather than biasing it towards a clade. A sensitivity analysis tested additional receptor keywords for *E. coli*, due to porins missing from the matrix, and returned unchanged nearest neighbors (*n* = 3) even though the feature vector was altered (S1 Text, S3 Table). Future versions could use ortholog protein clusters as features or further curated receptor vocabularies.

### Nearest-Neighbor Host Identification

PhagePickr receives a target bacterial species as input and queries NCBI for receptor data if the target is absent from the built-in dataset. Alternatively, users can supply receptor annotations through an optional argument. Without receptor annotations from the built-in dataset, NCBI, or user input, PhagePickr cannot generate biologically supported cocktails for the target species. When receptor annotations are available, the tool uses a nearest-neighbor algorithm to identify bacteria with similar receptor profiles. The scikit-learn v1.2.2 (46) NearestNeighbors subroutine acts on the Boolean receptor presence/absence vector (a row of the matrix *A*), and not the DNA or protein sequence of the host genome. The distance metric is available as an optional argument (Hamming, Jaccard, or cosine), with Hamming as the default. All reported cocktails were generated using Hamming distance. For two species with feature vectors *x* and *y*, the normalized Hamming distance is the fraction of differing receptor features, which is:

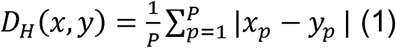

The numerator counts the mismatches between the Boolean vectors of two species, so *D_H_* is 0 for an identical configuration of receptor proteins and increases for each receptor gained or lost between species. PhagePickr then predicts cross-infectivity based on phenotypic similarity suggested by similar receptor profiles. Because *D_H_* is normalized over all *P* features and some species have few of them, the distance between two species cannot surpass the sum of their feature counts over *P*. Hence, *D_H_* reflects annotation depth in addition to receptor divergence and is used to rank neighbors rather than to compare distances across targets.

PhagePickr returns the *n* species with the smallest distance under the selected metric (*D_H_* for the default Hamming metric), which in default mode includes the target itself. The parameter *n* controls the number of neighbors and does not impose a minimum similarity to the target. For instance, using the Hamming distance, the nearest neighbors for *E. faecium* are *Staphylococcus pseudintermedius* (*D_H_*=0.0095) and *Clostridium perfringens* (*D_H_*=0.0102), corresponding to 28 and 30 differing features out of 2,951. However, *E. faecium* has 36 populated features, of which each neighbor shares 10. The small normalized Hamming distances therefore reflect shared absences rather than a substantial overlap in populated features. A similarity threshold could be implemented in future versions to allow the number of selected neighbors to vary. Upon identification of the nearest neighbors, each host contributes its registered phages to the candidate pool on which cocktail selection is performed.

Users of PhagePickr can keep the target species from contributing to the phage candidate pool, enabling exploratory phage discovery and comparison between validated and novel candidates (S1 Table). At present, this feature is limited to species listed in the built-in bacterial receptor dataset.

### Phage Selection

Phage selection is performed separately for each of the *n* hosts rather than over the entire pool of candidate hosts, as phages from different hosts may be too divergent to align meaningfully. For each host with more than two registered phages, the phage genomes are aligned with MAFFT (Multiple Alignment using Fast Fourier Transform) v7.526 (47) with automatic strategy selection (--auto) and an optional argument to set the number of threads, and a pairwise distance matrix *G* among potential phages is computed using BioPython’s identity distance. For a pair of aligned phages, *G_ij_* is the fraction of mismatched alignment columns including gaps and thus measures genomic divergence. A distance of 0 indicates identical genomes and larger values imply greater divergence. We define the most divergent phages as the pair with the largest *G_ij_*.

The phage pairs with the *k* highest distances are extracted using a size *k* min-heap evaluated across all pairwise distances, retaining the original phage pair indices (with a maximum of 2*k* phages per host). Hosts with two or fewer available phages do not require aligning and contribute all of them to the proposed phage cocktail. In the built-in dataset, 100 of the 200 receptor-matrix host entries (50%) have two or fewer registered phage accessions: 78 have one, and 22 have two. These reflect phage-host association database records rather than confirmed experimental infectivity against the target. Hosts without phage records are not included in the built-in dataset by design. The final cocktail then consists of the aggregation of selected phages across all *n* hosts. Fig 2 summarizes the basic process logic PhagePickr uses to propose the resulting phage cocktail.

**Fig 2.**
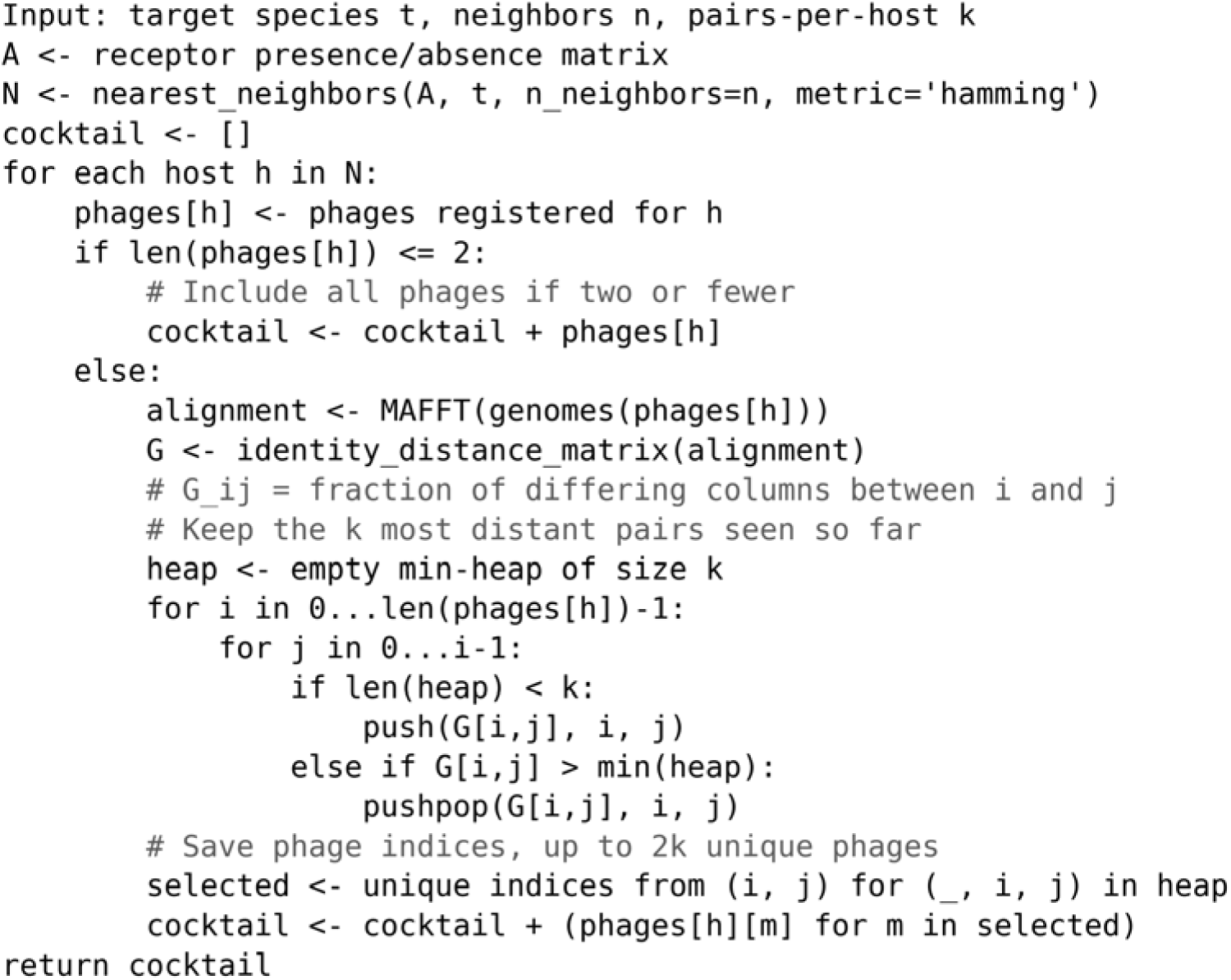
Pseudocode outlining the basic workflow for phage selection in PhagePickr. Once hosts with the most similar receptor profiles are identified, the workflow identifies phage pairs based on availability, or, if more than two are identified in the dataset, maximizes genomic divergence among phages.

The user-specified parameter *n* determines the breadth of the cocktail by governing the number of similar hosts contributing to the phage candidate pool, and the parameter *k* controls depth (the maximum number of highest divergence pairs retained per host). Moreover, *k* acts per host and is not a cap on cocktail size: each of the *n* hosts contributes at least one and at most 2*k* phages, so cocktail size ranges from *n* to 2*nk*. For instance, with defaults *n* = 3 and *k* = 1, each of the three hosts (including the target species) contributes its single most divergent pair, or its full set when two or fewer phages are available, resulting in final cocktails of three to six phages. The per-host independent alignment and selection prevent comparing extremely distant phage families and ensure each host neighbor is represented. This approach adapts the final candidate cocktail to bacteria with similar receptor configurations to the target. If these genomically diverse phages bind to different surface proteins, they could increase the likelihood of successful infection and make simultaneous resistance harder to evolve (28).

### Cocktail Diversity Scores

We examined the effect of the selection step on cocktail diversity using default parameter values (*n* = 3, *k* = 1). For each target, we identified its selection hosts (those with more than two registered phages); hosts with two or fewer contribute all their phages without a selection step and therefore have no most divergent pair to score. A target’s cocktail diversity score is the average maximum pairwise identity distance among phages from *H* selection hosts (the most divergent phage pair for *k* = 1):

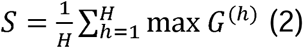

where *H* is the number of selection hosts, *G^(h)^* is the identity distance matrix for host *h*’s aligned phages, and the maximum is taken over all distinct phage pairs within that host. For comparison, the score was computed for random phage pairs per selection host, using the same per-host distances and across-host averaging, repeated 10,000 times to characterize the distribution of cocktail diversity for random controls. Species with a single selection host have a cocktail score that is identical to that host’s value rather than an average across hosts, and are shown for reference only.

### Lifestyle Prediction

An experimental feature to predict the lifestyle of phages in a cocktail using BACPHLIP v0.9.6 (48) is available as an optional argument. This feature is disabled by default and is not part of PhagePickr’s main workflow. Users can either obtain lifestyle predictions for the final PhagePickr cocktail, which just provides predicted annotations and leaves the cocktail unchanged, or apply a pre-selection filter that excludes all temperate phages from the candidate pool, generating exclusively lytic cocktails as predicted by BACPHLIP. This is not integrated within the main PhagePickr workflow and remains an experimental component. Hence, all reported cocktails were generated with this feature disabled.

### Software Implementation

PhagePickr is available on UNIX-based systems (MacOS, Linux) and Windows through WSL. Dependencies are managed via a Conda environment for reproducibility and ease of installation. Details on installation and usage are available on GitHub. To produce the cocktails against ESKAPE, PhagePickr was executed on the Bioinformatics Research Center Cluster at North Carolina State University with a maximum memory allocation of 32GB of RAM per node.

### Application to ESKAPE Pathogens

The tool was tested on the ESKAPE pathogens (*Enterococcus faecium*, *Staphylococcus aureus*, *Klebsiella pneumoniae*, *Acinetobacter baumannii*, *Pseudomonas aeruginosa*, *Enterobacter* spp.). These species were selected as a case study for resistance-aware cocktail design due to their clinical importance and well-documented antimicrobial resistance. The tool was also tested on two additional, less well-studied pathogens (*P. multocida* and *B. pseudomallei*).

## Results

The neighboring hosts for each target species were identified by a nearest-neighbor algorithm. These hosts supplied the candidate phages later selected into the cocktails (Table 1, S1 Table).

**Table 1.**
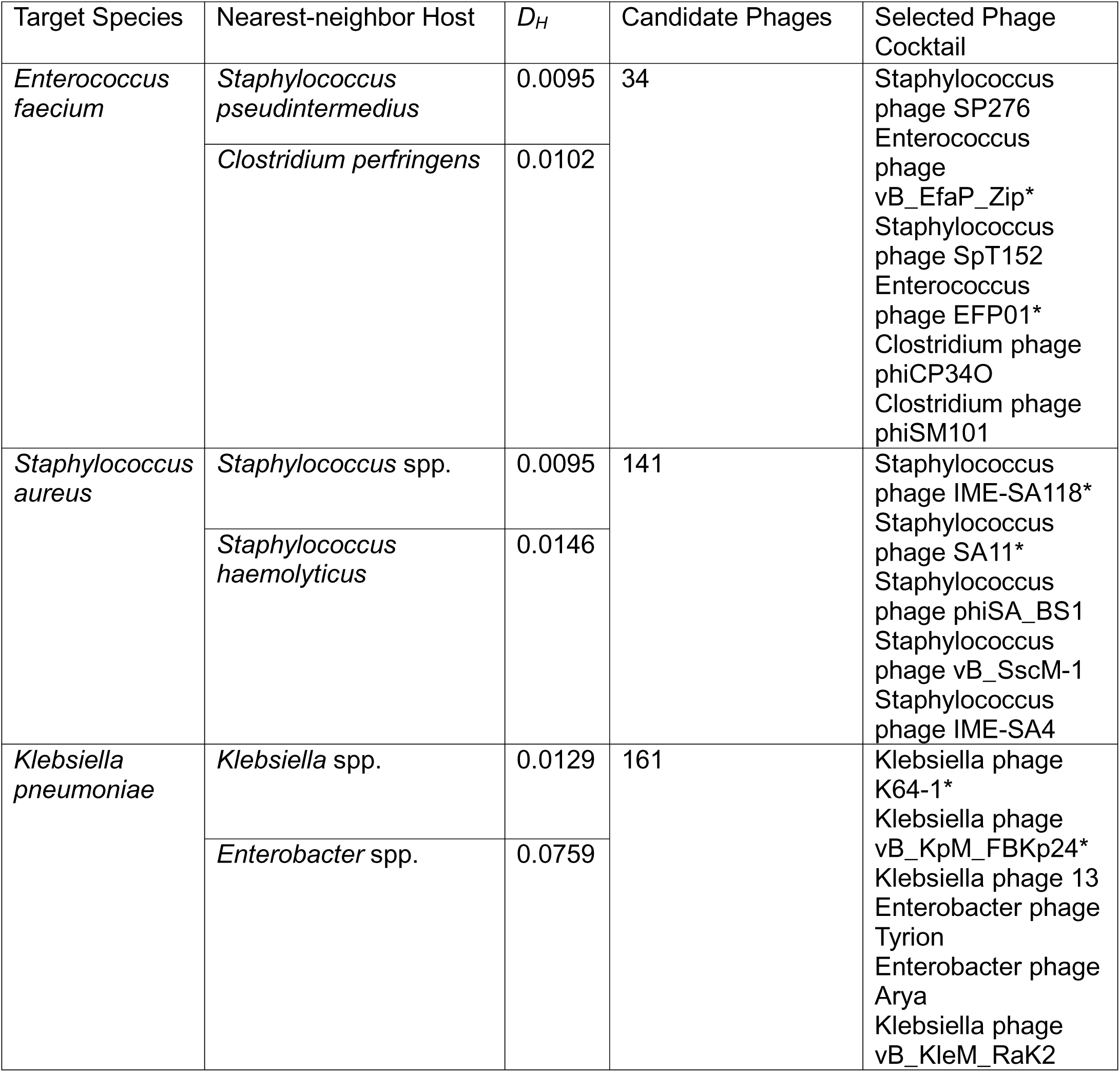
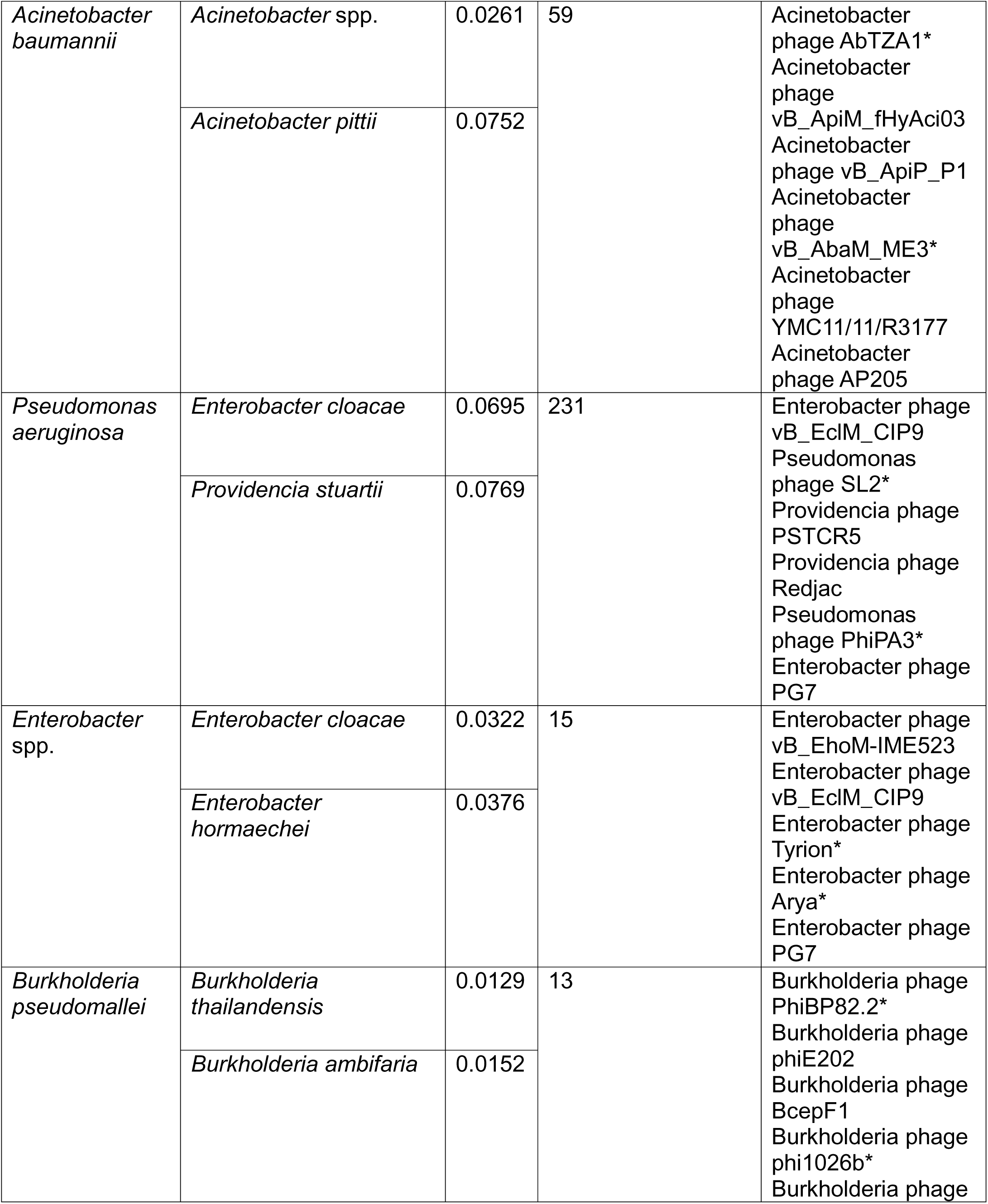
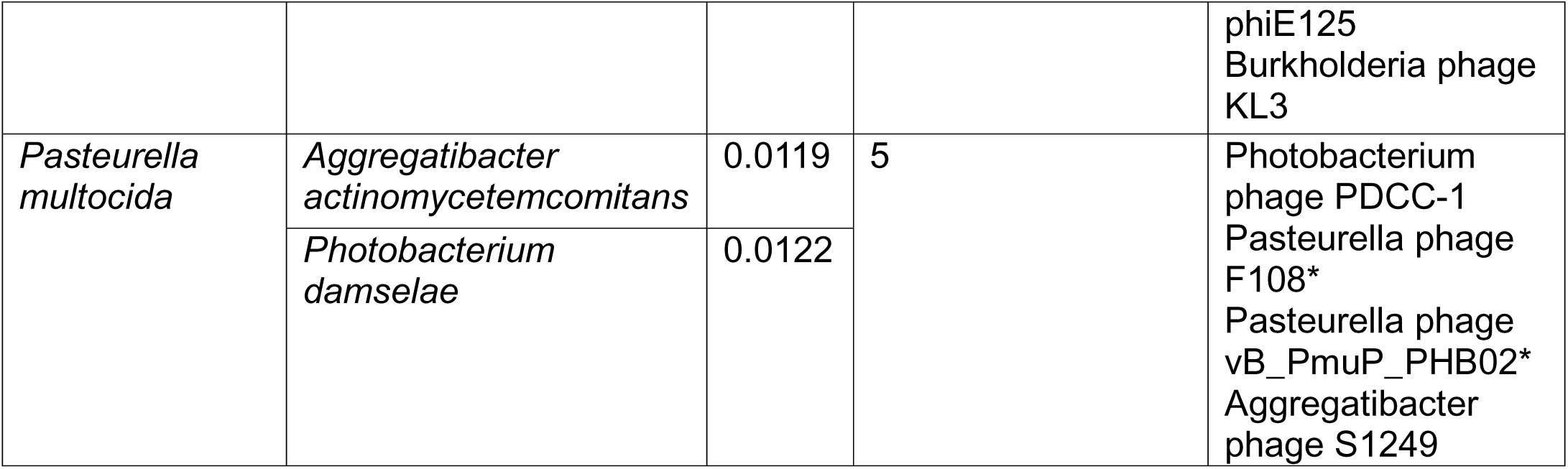
Phage cocktail composition and nearest-neighbor hosts for ESKAPE and understudied pathogens generated by PhagePickr (*n* = 3, *k* = 1). Here *n* is the number of phenotypically similar host species retrieved by the nearest-neighbor search. In default mode, the target species is counted among these *n*, thus the candidate pool spans the target and its *n*-1 closest neighbors. *k* is the maximum number of highest divergence phage pairs retained from each per-host alignment. *D_H_* is the normalized Hamming distance, or the fraction of differing receptor features between two species. The number of candidate phages is the union of the registered phages for the target and its *n*-1 nearest-neighbor hosts prior to diversity-based selection. The NCBI dataset includes data at both the species and genus level. Phage retrieval uses exact host labels, so a genus-level neighbor contributes only phage records labeled with that genus, rather than all records for its species. An asterisk (*) indicates phages previously reported to infect the target species.

We manually evaluated the predicted cocktails for the ESKAPE pathogens and two understudied pathogens (*Burkholderia pseudomallei* and *Pasteurella multocida*), with particular focus on their lysing ability, therapeutic potential, and host range (Table 2). Therapeutic potential was assessed relative to the bacterial species and strains examined by the cited studies, which may be the target, a neighbor, or another tested host. Evidence from other hosts does not demonstrate therapeutic potential against the target; the generated cocktails have not been experimentally validated.

**Table 2.** Summary of proposed cocktails generated by PhagePickr using default parameters (*n* = 3, *k* = 1). Therapeutic potential reflects literature support for individual candidates in the hosts examined by the cited studies and does not imply established efficacy of the cocktail. “Sourced from Target Genus” reports the fraction of phages in the final cocktail with hosts in the same genus as the target species. Cocktail size across the eight targets ranged from four to six phages. A host with more than two registered phages contributes its single most divergent pair, whereas a host with two or fewer contributes all its phages. We refer to a host with more than two phages as a selection host. The maximum cocktail size is therefore 2*nk* = 6, and a cocktail is smaller when one or more of the *n* hosts has only one registered phage. For details on individual phages within the cocktail, see the main text.

| Target Pathogen | Reported Lifestyle(s) | Therapeutic Potential | Reference | Sourced from Target Genus |
| --- | --- | --- | --- | --- |
| *E. faecium* | Lytic, temperate, unknown | Yes | (49,50) | 2/6 |
| *S. aureus* | Lytic, unknown | Yes | (51) | 5/5 |
| *K. pneumoniae* | Lytic, temperate, unknown | Yes | (52–56) | 4/6 |
| *A. baumannii* | Lytic | Yes | (57–61) | 6/6 |
| *P. aeruginosa* | Lytic, unknown | Yes | (62–66) | 2/6 |
| *Enterobacter* | Lytic, temperate, unknown | Yes | (55,56,64,65,67) | 5/5 |
| *B. pseudomallei* | Temperate | Validated prophages, potential for genetic engineering. | (68–70) | 6/6 |
| *P. multocida* | Lytic, temperate, unknown | Yes | (71–73) | 2/4 |

Notably, PhagePickr selected phages for *Enterococcus faecium* from distant genera, such as *Staphylococcus* and *Clostridium*, prioritizing shared receptor configuration rather than phylogenetic relatedness. Given the target’s limited receptor annotations, this cocktail is rather exploratory. Staphylococcus phages SP276 and SpT152 are likely temperate, but have therapeutic potential against *Staphylococcus pseudintermedius* given their similarity to phage VL4 (49). *Clostridium* phage phiSM101 encodes a powerful endolysin, effective against *Clostridium perfringens* (50). The identified candidate phages specific to *Enterococcus*, vB_EfaP_Zip and EFP01, lacked reliable data on infectivity and lifestyle.

By contrast, the cocktail for *Staphylococcus aureus* consisted of only *Staphylococcus* phages, as its closest pathogenic relatives also belong to this genus and are well studied. Among these, phage SA11 is a candidate for therapy with confirmed lytic activity against *S. aureus* ATCC 13301 (51), though the lifestyle and infectivity of other candidates, like IME-SA118, is largely unknown.

The *Klebsiella pneumoniae* cocktail included multiple phages with confirmed lytic activity from closely related genera *Klebsiella* and *Enterobacter*. For instance, Phages K64-1 and vB_KleM_RaK2 are both lytic and genetically similar (52). K64-1 encodes capsule depolymerase genes and has shown success in spot tests; while vB_KleM_RaK2 contains lysozymes and presents a host range likely specific to the *Klebsiella* genus (53). Moreover, phage vB_KpM_FBKp24 displays lytic activity in killing assays, but does not completely remove bacteria (54). Hence, its therapeutic use may depend on its inclusion in a cocktail. Phages Tyrion and Arya were also selected from an *Enterobacter* neighbor (Table 1).

The cocktail for *Acinetobacter baumannii* included only genus-specific phages: *Acinetobacter* phage AbTZA1, which encodes the endolysin *Ab*Lys1 and has direct killing potential (57), and confirmed lytic phage vB_ApiM_fHyAci03 (58). Further complementing the cocktail, phage vB_AbaM_ME3 produces lytic enzymes against *A. baumannii* (59), phage YMC11/11/R3177 has previously shown lytic potential (60), and phage AP205, a single-stranded RNA virus predicted to be lytic (61). Overall, the selected candidates have reported or predicted lytic potential, although the cocktail’s therapeutic efficacy is unverified.

The *Pseudomonas aeruginosa* cocktail evidenced the exploratory ability of PhagePickr due to the absence of close phylogenetic neighbors (Table 1). A diverse set of phages were selected, including those from *Providencia* and *Enterobacter*. The most propitious candidates PhagePickr identified were direct infectors of *P. aeruginosa*: phage SL2 has a large potential to cleanse planktonic cells but not against biofilms, infecting 13 of 24 *P. aeruginosa* strains (62). Similarly, phage PhiPA3 also has been shown to form plaques on different *P. aeruginosa* strains (63). The tool also proposed Enterobacter phages vB_EclM_CIP9, a likely lytic myovirus (64), and PG7, a *Straboviridae* phage potentially encoding lytic enzymes (65). However, it also included *Providencia stuartii* phages Redjac and PSTCR5. While the lifestyle of Redjac remains undetermined, PSTCR5 has been identified as lytic (66).

For *Enterobacter*, PhagePickr’s cocktail presented several phages with demonstrated efficacy. For instance, it selected Enterobacter phage vB_EhoM-IME523, shown to increase the survival rate of zebrafish infected with *E. hormaechei* under aeration conditions from 28.89% in controls not receiving the phage to 77.8% (67). As specified in the *P. aeruginosa* cocktail, Enterobacter phages vB_EclM_CIP9 and PG7 have lytic potential. Moreover, the suspected lysogenic Enterobacter phage Tyrion was included (55), as was phage Arya, which is highly syntenic to other lytic phages, but whose lifestyle remains unknown (56).

To assess PhagePickr’s ability to recognize candidate phages in underrepresented human and animal pathogens, we tested it on two less-studied species: *B. pseudomallei* and *P. multocida*. These two pathogens were selected because of the limited number of confirmed phages and the lower infection prevalence compared to ESKAPE. PhagePickr’s proposed *B. pseudomallei* cocktail included two validated prophages and four phages that infect other *Burkholderia* species. While these phages (BcepF1, phiE202, phiE125, KL3) display lysogenic behavior (68–70), they represent true infectors of close relatives to *B. pseudomallei* and remain viable candidates. In addition, these lysogenic phages can potentially be engineered to enter the lytic phase (74,75).

The cocktail for *P. multocida* included vB_PmuP_PHB02, which lyses specific capsular-type A strains with high efficacy (71) and the temperate phage F108 that lacks pathogenic elements (72). The tool identified *Photobacterium damselae* (previously known as *Pasteurella piscicida* (76)) as phenotypically similar to *P. multocida*, and included *Photobacterium* phage PDCC-1. Notably, *Photobacterium* is a phylogenetically distant genus, belonging to a different family and order. It also proposed *Aggregatibacter* phage S1249 which has shown lytic activity against *A. actinomycetemcomitans* (73), a member of the *Pasteurellaceae* family. Hence, the phylogenetic and phenotypic proximity between the host species makes phage S1249 a strong candidate against *P. multocida*.

We further summarize the cocktails reported in Table 2 by describing additional aspects of their composition. Of note, the selection step in PhagePickr retrieves the most divergent phage pair from each selection host. We scored each selected cocktail using the average maximum pairwise identity distance across its selection hosts and compared it against randomly assembled cocktails from the same host pools (Fig 3). Across the six targets with two or more selection hosts, selected cocktails had higher diversity scores than the random controls, most evidently for targets with large candidate phage pools. The difference is expected because PhagePickr maximizes the same genomic distance used to score the comparison. This therefore provides an implementation verification of the diversity maximization step, but does not serve as independent biological validation for receptor diversity or reduced cross-resistance.

**Fig 3.**
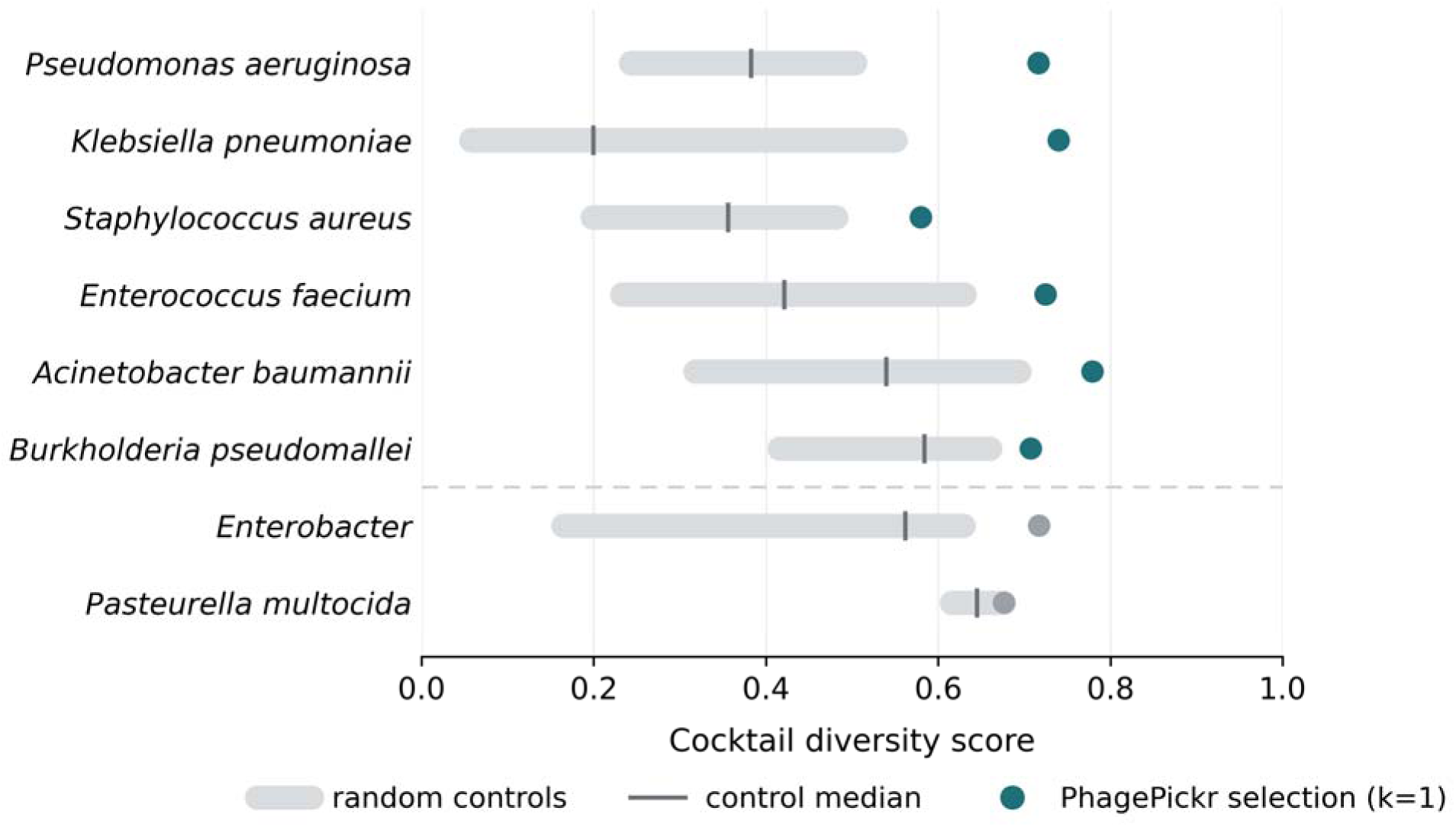
Effect of PhagePickr’s selection step on cocktail diversity. The cocktail score is the maximum pairwise identity distance among registered phages per host, averaged across a species’ selection hosts (hosts with more than two registered phages). The dot is the score for PhagePickr’s selection (*n* = 3, *k* = 1). The band spans the 5th-95th percentile of random controls (10,000 replicates, one random pair per selection host). A dot right of the band indicates the selected cocktail is more diverse than random pairing from the same pools. *Enterobacter* and *P. multocida* (grey) have a single selection host and are shown for reference only. Species are ordered by the standardized difference between the score of PhagePickr’s selection and the random controls (largest first); scores are not comparable across species.

The computational performance of PhagePickr was evaluated on all tested targets using two key metrics: memory allocation and runtime (S2 Table). Peak memory remained low relative to our memory allocation (32 GB) across all targets (131-1,515 MB; S2 Table), with the maximum usage observed for *P. aeruginosa*’s cocktail. Conversely, runtime increased with the total length of phage sequences aligned (Fig 4), with the multiple sequence alignment (MSA) of phage genomes being the main bottleneck. Total alignment runtime for the eight tested targets ranged from under a minute (*P. multocida*) to 59 h (*K. pneumoniae*). All runtimes and peak memory values reported were obtained with default single-thread settings. Due to this bottleneck, PhagePickr has an optional argument that allows the use of multiple cores in MAFFT. Since receptor annotations for all ESKAPE pathogens were pre-loaded in the PhagePickr dataset, no remote NCBI requests were made, thereby preventing any runtime delays due to network restrictions or server limitations.

**Fig 4.**
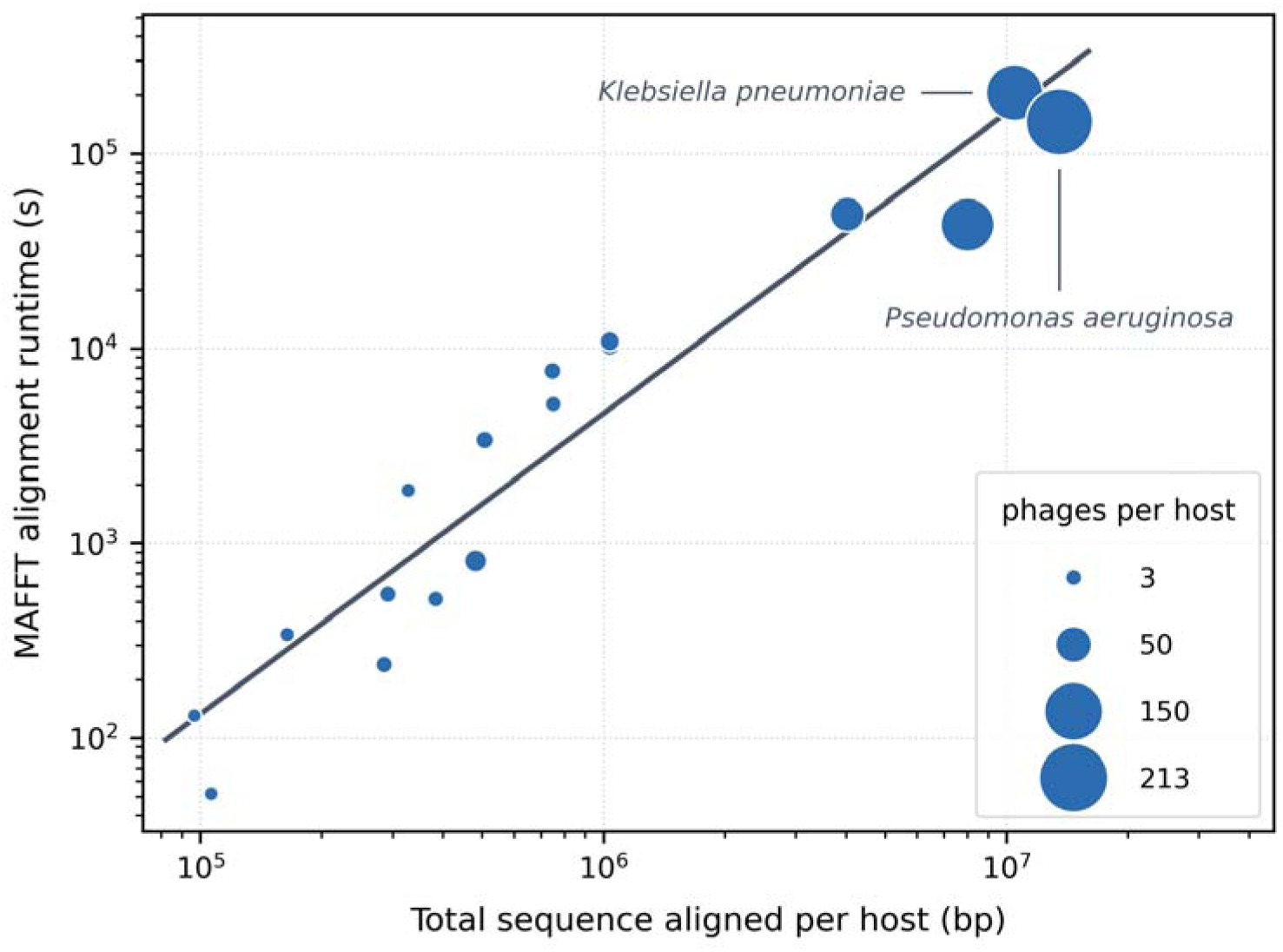
Scaling of per-host alignment runtime with sequence length. Each point is one alignment run of a selection host’s candidate phages across the eight target species (ESKAPE and understudied species, 17 runs involving 16 distinct selection hosts; two runs correspond to *E. cloacae*); the multiple sequence alignment (MSA) of each host’s candidate phages is the runtime bottleneck. The x-axis represents the total length of phage sequence aligned per host (bp); y-axis indicates MAFFT runtime (s) using one thread. Both the axes are on log scales, and point area is proportional to the number of phages aligned for that host. A straight-line fit on the log-scale axes has slope 1.54 (R^2^=0.92).

### Availability and Future Directions

The importance of phenotypic resistance and receptor-specific phage selection has been previously emphasized (77), supporting our strategy of selecting phages based on phenotypically similar bacteria by targeting their shared features to predict adsorption. Tools that predict phage hosts based on annotated receptor-binding proteins have also been proposed (78), motivating the framework described here, which follows an inverse approach that instead uses annotated receptor proteins in bacteria to predict phage infectors. While therapeutic value remains unverified, we evaluated PhagePickr’s success by whether it finds the phenotypically nearest-neighbor hosts by receptor profile and returns a complete cocktail composed of the most genomically divergent phage pairs from each selection host. Accordingly, PhagePickr successfully generated cocktails for a range of pathogens (including relatively understudied *P. multocida* and *B. pseudomallei*). Across all eight targets the pipeline returned a complete cocktail of four to six phages drawn from nearest-neighbor hosts, and every cocktail except that for *B. pseudomallei* contained at least one phage with reported lytic activity against the target or neighbor hosts (Table 2).

While PhagePickr’s cocktails alone could be tested for therapeutic utility, they can also be complemented with antibiotics for increased efficacy. Due to their different mechanisms of action, the simultaneous use of phages and antibiotics reduces the probability of resistance to either emerging permanently (9). Past studies have shown the potential synergistic effects of using phages and antimicrobial peptides (AMPs) against multi-drug resistant bacteria via lysis through host infection or cell membrane disruption (79). Similarly, it has been shown that the host can be rendered vulnerable to other phages once it develops resistance to one phage (80). Therefore, diversity among phages in PhagePickr cocktails is maximized to increase the opportunity for collateral sensitivity, under the hypothesis that divergent phages may bind different receptors.

Further, our receptor-based identification of phages has the potential to identify ways to re-sensitize bacteria to antibiotics. For instance, including phages that bind to efflux pumps, a common antibiotic resistance mechanism, could result in altering the pump, in turn leading to increased antibiotic susceptibility (31). This evolutionary trade-off suggests a critical synergy that PhagePickr cocktails could exploit downstream. By adding an optional layer of receptor selection on functional annotation after the host neighbors are found, the phages could target receptors related to antibiotic-resistance. These cocktails would have the potential to steer resistance evolution in hosts toward collateral sensitivity, which could be confirmed with in vitro testing of phages used in conjunction with antibiotics (81). As antibiotic-sensitive receptor targets become better characterized for more species, incorporating such a layer can be integrated into future iterations of PhagePickr.

A critical finding across several tested pathogens was the scarcity of functional annotation of phages. Many of the phages selected by PhagePickr have unclear therapeutic effects, due to a lack of published studies on infectivity, lifestyle, and host range. This was evident in the cocktails for *E. faecium* and *S. aureus*. Although some candidate phages were suspected lytic, most have not been tested in vitro, let alone in clinical trials. While we demonstrated the tool’s capacity to propose phages based on host receptor features, their therapeutic potential remains uncertain. Given that the purpose of this tool is to propose phage candidates to be characterized computationally, evaluated in the laboratory, and assayed in vivo, tools such as PhagePickr highlight a strong need for much more comprehensive genomic and phenotypic characterization followed by experimental validation of candidate phages.

Similar considerations apply to annotations for bacterial receptors. Such annotations must be dependable for PhagePickr to identify the biologically meaningful nearest neighbors. Because receptor annotation depth varies substantially across species, the cross-genus phage selection for species such as *E. faecium* is of a preliminary nature due to sparse annotations. As receptor annotations become richer, confidence in the selected nearest neighbors and the likelihood of therapeutic efficacy of the overall predicted cocktail increases. For instance, future characterization of receptor proteins in resistant bacterial strains may enable the exploration of strain-specific cocktails. Using PhagePickr to generate cocktails based on across-strain (rather than, or in addition to, across-species) receptor comparisons could prove especially promising for resistance-aware phage cocktail design, as the resulting phages from related strains may prove quite virulent against novel hosts and manage coinfections involving multiple strains. We believe the availability of tools like PhagePickr could therefore spur improved strain-level receptor annotation data availability.

Thus, given existing data limitations PhagePickr remains a hypothesis-generating tool. Our computational approach can help these experimental methods realize their full potential by rapidly suggesting a large set of candidate phage assays that could be performed in vitro against strains of the target species, prior to clinical applications: host-range spot tests, efficiency of plating, killing curve analyses, and adsorption assays. The tool’s applicability for different species could also be evaluated experimentally, which would require testing the efficacy of phage cocktails identified by PhagePickr (from receptor-similar hosts) against control phage cocktails (e.g., from receptor-dissimilar hosts).

A further consideration is that phage therapy still presents important safety challenges. Interactions with other drugs, transfer of undesired genetic material to pathogens or microbiome, phage antibody production, and the release of endotoxins in Gram-negative bacteria can all interfere with the efficacy of phage therapy or lead to adverse health outcomes (82,83). In addition, it is recommended that therapeutic phages do not have integrases, virulence factors, or genes associated with antibiotic resistance (84). As phage annotations and characterizations improve, screening phage genomes for such genes downstream could be incorporated into future iterations of the tool.

How best to enable PhagePickr to distinguish lytic from temperate phages at present remains an open question in our view. The clearance of pathogenic bacteria generally involves lysis, and phages that keep their hosts alive might not be immediately useful if the objective is to heal a clinical infection. However, some users may wish to retain lysogenic phages, as such phages can still have natural antimicrobial activity without altering bacterial pathogenicity (85), be genetically engineered for lytic function (74,75), or act as bacteriostatic agents disrupting essential gene pathways (86).

Even for users wishing to screen lysogenic phages, there is at present no single prevailing method allowing for the extraction of lifestyle features from database entries, since this requires previous extensive bioinformatic and experimental characterization of phages. As a potential solution, however, there are several bioinformatic tools that allow for phage lifestyle prediction from genome sequence. These include BACPHLIP (48), DeePhage (87), PhagePred (88), and PHACTS (89), which use a range of methods to classify phage lifestyles: convolutional neural networks, alignment-free methods, or Random Forest models. Such computational tools could serve as a complement to the genome screening described above to filter lysogenic candidate phages proposed by PhagePickr. As a proof of principle, we show that a lifestyle cross-check with existing methods is a feasible downstream step (S2 Text, S4 Table). However, we have elected to keep this as an experimental feature via an optional argument for now (see Design and Implementation) as no single lifestyle predictor method is currently standard, and available tools differ considerably in approach, training data, and maintenance status.

Nearest-neighbor selection is also sensitive to the choice of metric for receptor similarity. Comparing normalized Hamming and Jaccard distances at *n* = 3 changed at least one of two selected neighbors, excluding the target species itself, for five of eight targets. These changes included both neighbors of *E. faecium* and *P. aeruginosa* (S3 Text, S5 Table). However, this comparison does not establish which metric is best at identifying biologically appropriate hosts.

The tool’s exploratory feature can represent a limitation if too many neighbors are considered in the analysis, particularly if the species has no close relatives in the dataset. The reliance on receptor similarity can lead to the selection of phages from relatively distant genera (as with *E. faecium*, where annotations are sparse, and *P. aeruginosa*, where no close neighbor is available). Consequently, the probability of phage adsorption and infection may decrease with increased phylogenetic distances of neighbors included. Nonetheless, the algorithm can be refined to limit the number of relatives used and keep phage inclusion restricted, thereby potentially improving cocktail efficacy even if at the cost of phage diversity. Finally, we add that even if the phages proposed by PhagePickr do result in a mismatch with the host, they still remain promising candidates for genetic engineering and recombineering, due to the similarity between host receptor profiles.

PhagePickr designs cocktails based on the likelihood of adsorption via shared receptors. However, phage adsorption is only the first step for a successful infection and killing of the host. Other factors that can influence the effectiveness of phage therapy include infection duration, phage burst size, and virion durability (90). Due to limited data on these criteria for most sequenced phages, these factors are not considered in the tool’s current iteration but are candidates for future inclusion as selection criteria.

We found PhagePickr’s computational performance to be determined by total sequence length (Fig 4), as it directly impacts the MSA step. As expected, the MSA of entire phage genomes, which we use to maximize diversity, is the rate-limiting step of the workflow. Although sequence imperfectly predicts function, a cocktail with divergent phages may reduce the risk of cross-resistance from similar infection pathways (91). While PhagePickr does offer an alignment-free alternative (random phage selection) to accommodate computational resource and time constraints, maximum cocktail diversity is preferred to hinder evolutionary resistance.

Despite these potential limitations, PhagePickr nevertheless provides a much-needed framework to address the emergence of phage resistance in bacteria. To our knowledge, this is the first attempt at an automated, systematic and reproducible receptor-based phage cocktail prediction tool designed specifically to circumvent the evolution of anti-phage resistance. Moreover, many of the potential limitations are due to data availability. Thus, we anticipate our tool’s utility and predictive potential will scale considerably as more phage and bacterial receptor data become available. In addition to the ability to generate cocktails from cross-strain comparisons discussed above, future directions upon the framework described here include tool extensions such as validation of the existing optional lifestyle filter to exclude temperate phages and an optional antibiotic-resistance-related receptor downselection mode to steer collateral sensitivity; and downstream applications of the output like pairing cocktails with antibiotics and AMPs as a combination strategy. We expect these additions to further improve PhagePickr’s ability to design resistance-aware phage cocktails.

The source code for PhagePickr is publicly available on GitHub (https://github.com/Alebraco/phagepickr) under the MIT license.

## Supporting Information

S1 Table. Exploratory cocktail composition for ESKAPE and understudied pathogens (*n* = 3, *k* = 1).

S2 Table. Computational performance metrics of MAFFT phage alignments under default and exploratory mode for ESKAPE and understudied pathogens (*n* = 3, *k* = 1).

S3 Table. Nearest-neighbor robustness for *Escherichia coli* under an expanded receptor search.

S4 Table. Comparison of automated phage-lifestyle predictions against literature-based classifications for 40 phages across eight phage cocktails.

S5 Table. Comparison of non-target nearest-neighbor hosts under normalized Hamming and Jaccard distances (*n* = 3, default mode).

S1 Text. Sensitivity of nearest-neighbor selection to receptor keyword search.

S2 Text. Phage lifestyle classification on selected cocktails.

S3 Text. Sensitivity of nearest-neighbor selection to receptor similarity metric.

