## Supporting Information for "PhagePickr: A bacteria-centric computational tool for designing resistance-aware phage cocktails"

**S1 Table. Exploratory cocktail composition for ESKAPE and understudied pathogens (*n* = 3, *k* = 1).** Exploratory cocktails were generated using the --explore_only argument in PhagePickr, in which the target species still defines the nearest-neighbor search but contributes no phages. The cocktail therefore draws phages from its *n* non-target neighbors. See Table 1 for definitions of *n* and *k* (in exploratory mode, the target is not counted among the *n*).

| Target Species | Exploratory Cocktail |
| --- | --- |
| *Enterococcus faecium* | Staphylococcus phage SP276  Staphylococcus phage SpT152  Staphylococcus phage IME-SA4  Clostridium phage phiCP34O  Clostridium phage phiSM101 |
| *Staphylococcus aureus* | Staphylococcus phage phiSA_BS1  Staphylococcus phage vB_SscM-1  Staphylococcus phage IME-SA4  Clostridium phage phiCP34O  Clostridium phage phiSM101 |
| *Klebsiella pneumoniae* | Klebsiella phage 13  Klebsiella phage KOX1  Enterobacter phage Tyrion  Enterobacter phage Arya  Klebsiella phage PKO111  Klebsiella phage vB_KleM_RaK2 |
| *Acinetobacter baumannii* | Enterobacter phage vB_EclM_CIP9  Acinetobacter phage vB_ApiM_fHyAci03  Acinetobacter phage vB_ApiP_P1  Acinetobacter phage YMC11/11/R3177  Enterobacter phage PG7  Acinetobacter phage AP205 |
| *Pseudomonas aeruginosa* | Providencia phage PSTCR5  Enterobacter phage vB_EclM_CIP9  Enterobacter phage phiEap-3  Enterobacter phage phiEap-1  Enterobacter phage PG7  Providencia phage Redjac |
| *Enterobacter* | Enterobacter phage vB_EhoM-IME523  Enterobacter phage vB_EclM_CIP9  Enterobacter phage phiEap-3  Enterobacter phage phiEap-1  Enterobacter phage PG7 |
| *Burkholderia pseudomallei* | Burkholderia phage phiE202  Burkholderia phage BcepF1  Burkholderia phage phiE125  Burkholderia phage KL3  Burkholderia phage KS9 |
| *Pasteurella multocida* | Photobacterium phage PDCC-1  Proteus phage phiP4-3  Aggregatibacter phage S1249 |

**S2 Table. Computational performance metrics of MAFFT phage alignments under default and exploratory mode for ESKAPE and understudied pathogens (*n* = 3, *k* = 1).** Cocktail composition is available for default (Table 1) and exploratory modes (S1 Table). See Table 1 for definitions of *n* and *k*. All alignments were run single-threaded. No alignment was necessary for the exploratory cocktail of *P. multocida* as its neighbors have one phage each.

| **Target Species** | **Default mode** | | **Exploratory mode** | |
| --- | --- | --- | --- | --- |
|  | Runtime | Peak Mem | Runtime | Peak Mem |
| *Enterococcus faecium* | 1 h 13 min | 270 MB | 17 min | 270 MB |
| *Staphylococcus aureus* | 13 h 25 min | 796 MB | 1 h 39 min | 270 MB |
| *Klebsiella pneumoniae* | 59 h 12 min | 1324 MB | 2 h 27 min | 380 MB |
| *Acinetobacter baumannii* | 14 h 06 min | 494 MB | 3 h 29 min | 312 MB |
| *Pseudomonas aeruginosa* | 43 h 37 min | 1515 MB | 3 h 17 min | 334 MB |
| *Enterobacter* | 2 h 52 min | 312 MB | 3 h | 312 MB |
| *Burkholderia pseudomallei* | 14 min 46 s | 270 MB | 5 min 40 s | 270 MB |
| *Pasteurella multocida* | 52 s | 131 MB | no alignment | n/a |

**S3 Table. Nearest-neighbor robustness for *Escherichia coli* under an expanded receptor search.** Nearest neighbors (*n* = 3) and Hamming distances (*D_H_*) under the original receptor search, the broadened search (projected vector; 2,951 original columns), and the broadened search with all columns (extended vector; 2,951 + 1,129 new columns). Distances for the extended vector are inflated because the added columns are present only for *E. coli*; the projected vector remains comparable to the original in column space. Within the 2,951 shared columns, the projected vector differs from the original at 90 positions, which explains the shifts in *D_H_*.

| Rank | Original search | *D_H_* | Expanded keywords, projected vector | *D_H_* | Expanded keywords, extended vector | *D_H_* |
| --- | --- | --- | --- | --- | --- | --- |
| 1 | *Escherichia spp.* | 0.0017 | *Escherichia spp.* | 0.0322 | *Escherichia spp.* | 0.3000 |
| 2 | *Shigella* | 0.1362 | *Shigella* | 0.1532 | *Shigella* | 0.3875 |
| 3 | *Shigella sonnei* | 0.1423 | *Shigella sonnei* | 0.1586 | *Shigella sonnei* | 0.3914 |

**S4 Table. Comparison of automated phage-lifestyle predictions against literature-based classifications for 40 phages across eight phage cocktails.** Predictions were generated with BACPHLIP v0.9.6 as an example among many available lifestyle classifiers (e.g. DeePhage, PhagePred, PHACTS). P(Virulent) and P(Temperate) are probabilities assigned by BACPHLIP for each lifestyle; the final prediction is the higher of the two values. The agreement column compares BACPHLIP’s prediction with the literature lifestyle (“—” indicates undetermined lifestyle in the literature). Of 30 phages with a defined literature lifestyle, 26 agreed with the prediction. This function is disabled by default and not part of the main workflow described in the main text.

| **Phage** | **Target** | **Literature Lifestyle** | **P(Virulent)** | **P(Temperate)** | **BACPHLIP prediction** | **Agreement** |
| --- | --- | --- | --- | --- | --- | --- |
| Staphylococcus phage SP276 | *Enterococcus faecium* | temperate | 0.00 | 1.00 | temperate | match |
| Enterococcus phage vB_EfaP_Zip |  | unknown | 0.98 | 0.02 | lytic | — |
| Staphylococcus phage SpT152 |  | temperate | 0.00 | 1.00 | temperate | match |
| Enterococcus phage EFP01 |  | unknown | 0.91 | 0.09 | lytic | — |
| Clostridium phage phiCP34O |  | lytic | 0.96 | 0.04 | lytic | match |
| Clostridium phage phiSM101 |  | temperate | 0.67 | 0.33 | lytic | mismatch |
| Staphylococcus phage IME-SA118 | *Staphylococcus aureus* | unknown | 0.71 | 0.29 | lytic | — |
| Staphylococcus phage SA11 |  | lytic | 0.85 | 0.15 | lytic | match |
| Staphylococcus phage phiSA_BS1 |  | unknown | 0.85 | 0.15 | lytic | — |
| Staphylococcus phage vB_SscM-1 |  | unknown | 0.91 | 0.09 | lytic | — |
| Staphylococcus phage IME-SA4 |  | unknown | 0.01 | 0.99 | temperate | — |
| Klebsiella phage K64-1 | *Klebsiella pneumoniae* | lytic | 0.99 | 0.01 | lytic | match |
| Klebsiella phage vB_KpM_FBKp24 |  | lytic | 0.70 | 0.30 | lytic | match |
| Klebsiella phage 13 |  | unknown | 0.95 | 0.05 | lytic | — |
| Klebsiella phage vB_KleM_RaK2 |  | lytic | 1.00 | 0.00 | lytic | match |
| Enterobacter phage Tyrion | *Klebsiella pneumoniae; Enterobacter spp.* | temperate | 0.04 | 0.96 | temperate | match |
| Enterobacter phage Arya | *Klebsiella pneumoniae; Enterobacter spp.* | unknown | 0.75 | 0.25 | lytic | — |
| Acinetobacter phage AbTZA1 | *Acinetobacter baumannii* | lytic | 1.00 | 0.00 | lytic | match |
| Acinetobacter phage vB_ApiM_fHyAci03 |  | lytic | 0.93 | 0.07 | lytic | match |
| Acinetobacter phage vB_ApiP_P1 |  | lytic | 0.81 | 0.19 | lytic | match |
| Acinetobacter phage vB_AbaM_ME3 |  | lytic | 0.96 | 0.04 | lytic | match |
| Acinetobacter phage YMC11/11/R3177 |  | lytic | 0.07 | 0.93 | temperate | mismatch |
| Acinetobacter phage AP205 |  | lytic | 0.81 | 0.19 | lytic | match |
| Pseudomonas phage SL2 | *Pseudomonas aeruginosa* | lytic | 0.94 | 0.06 | lytic | match |
| Providencia phage PSTCR5 |  | lytic | 0.89 | 0.11 | lytic | match |
| Providencia phage Redjac |  | unknown | 1.00 | 0.00 | lytic | — |
| Pseudomonas phage PhiPA3 |  | lytic | 0.84 | 0.16 | lytic | match |
| Enterobacter phage vB_EclM_CIP9 | *Pseudomonas aeruginosa; Enterobacter spp.* | lytic | 0.89 | 0.11 | lytic | match |
| Enterobacter phage PG7 | *Pseudomonas aeruginosa; Enterobacter spp.* | lytic | 1.00 | 0.00 | lytic | match |
| Enterobacter phage vB_EhoM-IME523 | *Enterobacter spp.* | lytic | 0.96 | 0.04 | lytic | match |
| Burkholderia phage PhiBP82.2 | *Burkholderia pseudomallei* | temperate | 0.00 | 1.00 | temperate | match |
| Burkholderia phage phiE202 |  | temperate | 0.01 | 0.99 | temperate | match |
| Burkholderia phage BcepF1 |  | temperate | 0.72 | 0.28 | lytic | mismatch |
| Burkholderia phage phi1026b |  | temperate | 0.00 | 1.00 | temperate | match |
| Burkholderia phage phiE125 |  | temperate | 0.01 | 0.99 | temperate | match |
| Burkholderia phage KL3 |  | temperate | 0.00 | 1.00 | temperate | match |
| Photobacterium phage PDCC-1 | *Pasteurella multocida* | unknown | 0.99 | 0.01 | lytic | — |
| Pasteurella phage F108 |  | temperate | 0.00 | 1.00 | temperate | match |
| Pasteurella phage vB_PmuP_PHB02 |  | lytic | 1.00 | 0.00 | lytic | match |
| Aggregatibacter phage S1249 |  | lytic | 0.01 | 0.99 | temperate | mismatch |

**S5 Table. Comparison of non-target nearest-neighbor hosts under normalized Hamming and Jaccard distances (*n* = 3, default mode).** Selected hosts are listed in increasing distance. Overlap counts shared host identities regardless of order, excluding the target. All host profiles were populated. Genus names refer to genus-level entries in the receptor matrix. *Ties occurred during nearest-neighbor selection. Only the recorded selections are shown. Alternative hosts at the same distance retain the reported overlap for *E. faecium* but can reduce it for *P. multocida*.

| **Target** | Hamming neighbors (*D_H_*) | Jaccard neighbors (*D_J_*) | Overlap |
| --- | --- | --- | --- |
| *E. faecium** | *Staphylococcus pseudintermedius* (0.0095)  *Clostridium perfringens* (0.0102) | *Enterococcus* (0.4930)  *Enterococcus faecalis* (0.6780) | 0/2 |
| *S. aureus* | *Staphylococcus* (0.0095)  *Staphylococcus haemolyticus* (0.0146) | *Staphylococcus* (0.3373)  *Staphylococcus epidermidis* (0.7761) | 1/2 |
| *K. pneumoniae* | *Klebsiella* (0.0129)  *Enterobacter* (0.0759) | *Klebsiella* (0.1070)  *Enterobacter* (0.5803) | 2/2 |
| *A. baumannii* | *Acinetobacter* (0.0261)  *Acinetobacter pittii* (0.0752) | *Acinetobacter* (0.2312)  *Pseudomonas aeruginosa* (0.6944) | 1/2 |
| *P. aeruginosa* | *Enterobacter cloacae* (0.0695)  *Providencia stuartii* (0.0769) | *Pseudomonas* (0.5824)  *Enterobacter* (0.6630) | 0/2 |
| *Enterobacter* spp. | *Enterobacter cloacae* (0.0322)  *Enterobacter hormaechei* (0.0376) | *Enterobacter cloacae* (0.4113)  *Enterobacter hormaechei* (0.4805) | 2/2 |
| *B. pseudomallei* | *Burkholderia thailandensis* (0.0129)  *Burkholderia ambifaria* (0.0152) | *Burkholderia thailandensis* (0.6786)  *Burkholderia cepacia* (0.7273) | 1/2 |
| *P. multocida** | *Aggregatibacter actinomycetemcomitans* (0.0119)  *Photobacterium damselae* (0.0122) | *Aggregatibacter actinomycetemcomitans* (0.6863)  *Photobacterium damselae* (0.7347) | 2/2 |

**S1 Text. Sensitivity of nearest-neighbor selection to receptor keyword search.**

A sensitivity analysis was performed on *Escherichia coli* to test the robustness of the keywords used to retrieve receptor proteins. Important *E. coli* outer membrane porins OmpC and OmpF were missed by the “receptor[All fields]” keyword. We assessed whether an increase of breadth in our keyword search, explicitly including porins and outer membrane proteins, would change the top nearest neighbors for *E. coli*. If the nearest-neighbor set is stable for *E. coli*, which is vulnerable to missing annotations, it is unlikely to shift for less-annotated species.

The NCBI IPG retrieval for *E. coli* was executed using the following terms: Escherichia coli[ORGN] AND (receptor[All fields] OR porin[All fields] OR “outer membrane protein”[All fields]). This returned 1,728 unique protein titles, of which 1,129 were not in the original feature matrix. Among the new features were 17 OmpC and OmpF variants, not captured by the original keywords. We then recomputed the *E. coli* feature vector in two ways: projected onto the original vector of 2,951, and with 1,129 new columns (2,951 + 1,129 total) corresponding to the new proteins captured by the expanded keyword set. The nearest-neighbor algorithm (*n* = 3, metric = “hamming”) was applied to both, with *E. coli* excluded from the set (as in exploratory mode), so that all neighbors are different species.

The set of nearest neighbors (*n* = 3) and its ranking were unchanged for both applications (S3 Table). Within the original vector column space (2,951 features), the new projected *E. coli* vector differed from the original at 90 positions (60 features gained, 30 lost, 3.1% of vector). Features absent from the recomputed vector reflect changes in IPG entries since the feature matrix was built rather than keyword change, as the expanded search still contains the original terms. Thus, an altered feature vector recovered the same neighbors. We believe this is evidence that the neighbor selection is robust to the keyword choice in receptor search.

**S2 Text. Phage lifestyle classification on selected cocktails.**

A lifestyle classification downstream step was performed on the predicted phage cocktails. The selected phage genomes were passed to a lifestyle prediction tool to cross-check against the literature. Complete RefSeq genomes were retrieved for the 40 phages across the eight proposed cocktails and processed with BACPHLIP v0.9.6, which classifies genomes as lytic or temperate from the presence of temperate-associated protein domains. BACPHLIP was used for convenience as an example, but any tool that uses whole genome sequences could be used. Lifestyle predictions were compared with the literature for each phage, and phages for which the literature is inconsistent or unavailable were recorded as unknown.

BACPHLIP was able to classify all 40 phages. Ten phages with unknown lifestyle were classified, but these predictions cannot be compared to the literature due to lack of data. Notably, the automated predictor agreed with 26 out of 30 (87%) phages with a literature-defined lifestyle, including well-characterized prophages and virulent phages. However, it disagreed with four phages in both directions. It classified Clostridium phage phiSM101 and Burkholderia phage BcepF1 as lytic, whereas Acinetobacter phage YMC11/11/R3177 and Aggregatibacter phage S1249 were confidently deemed temperate phages, inconsistent with literature claims.

While useful, the automated classification of phage lifestyle still conflicted with the literature in both directions. Integrating lifestyle prediction within the main PhagePickr workflow could either admit temperate phages the filter fails to flag or exclude truly lytic ones, which undermines the purpose of using such tools. Because lifestyle prediction and temperate phage exclusion remain tool dependent and no method is currently standard, these are disabled by default and only available via the optional argument “--lifestyle” in PhagePickr.

**S3 Text. Sensitivity of nearest-neighbor selection to receptor similarity metric.**

We compared normalized Hamming and Jaccard distances using the same receptor matrix (200 hosts, 2,951 features) for the eight tested targets. All host profiles contained at least one populated feature. For two hosts with feature sets A and B, the Jaccard distance is the fraction of features present in either host that are not shared, which is:

$$D_{J}=1-\frac{\left| A\cap B \right|}{|A\cup B|}$$

This excludes shared absences and is defined when the union is nonzero. With *n* = 3 in default mode, we compared the two selected non-target neighbors under each metric. Top-n overlap was recorded as the number of shared host identities out of two, regardless of order. S5 Table reports the selected hosts, their distances, and overlap for each of the eight targets.
